# *In vitro* Cleavage Requirements and Specificities of Mycobacterial RNase E

**DOI:** 10.64898/2026.04.06.716707

**Authors:** Abigail R. Rapiejko, Manchi Reddy, James C. Sacchettini, Scarlet S. Shell

## Abstract

Regulation of RNA pools allows for adaptation to changing environments and stress, which is especially important in pathogenic bacteria such as *Mycobacterium tuberculosis*. RNA degradation is a significant contributor to RNA abundance, and Ribonuclease (RNase) E has a rate-limiting role in degradation of a majority of mycobacterial transcripts. However, many open questions remain about the RNA substrate requirements and specificities for efficient cleavage by mycobacterial RNase E. Here, using both *Mycolicibacterium smegmatis* and *M. tuberculosis* RNase E, we demonstrate that this enzyme is only active on substrates with a minimum length of approximately 27 nt. Furthermore, we show that mycobacterial RNase E prefers substrates with 5’ monophosphates rather than 5’ triphosphates, and that the positions of cleavage events within substrates are dictated by both sequence and distance from the RNA ends. Our results also suggest that RNase E may be affected by product inhibition. Finally, we show that *M. smegmatis* RNase E behaves similarly to *M. tuberculosis* RNase E, validating the use of this model organism for RNA degradation studies.

## INTRODUCTION

*Mycobacterium tuberculosis* remains a global public health threat with millions of reported cases and deaths each year [1]. An important enzyme likely regulated during physiologically relevant stress is RNase E. This enzyme is an endonuclease that has a rate-limiting role in bulk mRNA degradation in the nonpathogenic model mycobacterium *Mycolicibacterium smegmatis* [2] and is essential for growth in both *M. smegmatis* and *M. tuberculosis* [3, 4].

In *Escherichia coli*, the catalytic domain of RNase E contains an S1 subdomain for RNA binding, a 5’-end-sensing pocket, and DNase I-like and RNase H-like subdomains [5]. Multiple sequence alignments of *E. coli* RNase E with *M. tuberculosis* RNase E suggest that this catalytic domain architecture is conserved [6]. As a result, other similarities exist between *E. coli* and mycobacterial RNase E such as the requirement for magnesium in the active site [5, 7] and the use of zinc for the formation of tetramers (dimer of dimers) [5, 7, 8]. However, mycobacterial RNase E also displays many differences from its better-studied *E. coli* counterpart. Mycobacterial RNase E has two intrinsically disordered regions (IDRs) flanking the catalytic domain, unlike *E. coli* RNase E which has only a single IDR [6]that scaffolds a multiprotein complex called the degradosome. The degradosome is formed by stable interactions in *E. coli,* whereas mycobacterial degradosome is predicted to be transient due to the need for crosslinking to identify its components [4]. *E. coli* RNase E has a membrane binding domain and is therefore localized to the inner surface of the cell membrane [9–14], while mycobacterial RNase E is cytoplasmic [15]. Finally, mycobacterial RNase E preferentially cleaves preceding cytidine residues, consistent with mycobacteria being GC-rich organisms [2], whereas *E. coli* RNase E preferentially cleaves preceding uracil residues, consistent with it being a GC-balanced organism [10, 16]. These important differences motivate studying mycobacterial RNase E, as it is fundamentally different from the better studied *E. coli* version.

Unanswered questions remain surrounding RNA substrate preferences and specificities of cleavage site position for mycobacterial RNase E. Here we used *in vitro* RNA cleavage assays with substrates containing exactly one known RNase E cleavage site to investigate these preferences and specificities with both *M. tuberculosis* and *M. smegmatis* versions of the enzyme. We found that RNase E has a minimum substrate length requirement of approximately 27 nt for efficient cleavage, in contrast to a previous report [7], and we suggest that RNase preparation purity may have contributed to these discrepancies. Additionally, we demonstrate that both RNases E and J prefer to cleave RNA substrates with 5’ monophosphates to those with 5’ triphosphates. We also show that cleavage site proximity to the RNA ends is just as important as cleavage site sequence for dictating RNase E cleavage position. Finally, we suggest that RNase E may be influenced by product inhibition *in vitro*. These findings provide a framework for future investigations using RNase E *in vitro* and have important *in vivo* implications towards understanding how RNase E selects and degrades its substrates to control RNA metabolism in a globally important pathogen.

## MATERIALS AND METHODS

### Bacterial Strains and Culture Conditions

RNase E variants as described below were overexpressed from pET-based plasmids in *E. coli* BL21 (DE3). Lennox Luria-Bertani broth and solid media were used to grow *E. coli*. Antibiotic concentrations used for *E. coli* were 50 μg/mL kanamycin and 20 μg/mL chloramphenicol. *E. coli* NEB-5-alpha (New England Biolabs) was used for cloning. HiFi Assembly (New England Biolabs) was used to make all plasmids for this study.

### Overexpression and Purification of Recombinant *M. smegmatis* Proteins

RNase E variants were recombinantly expressed and purified using *E. coli* BL21(DE3) pLysS transformed with pET42-derived plasmids. Gene IDs are listed in Supplemental Table 1 and expression plasmid descriptions are provided in Supplemental Table 2. 2 L cultures were grown at 37°C and were shaken at 200 rpm to an OD_600 nm_ of 0.5-0.8. Cultures were induced with 1 mM IPTG at 28°C and were shaken at 200 rpm for 4 hours prior to harvest by centrifugation. Cell pellets were resuspended in 10 mL of buffer (20 mM tris-HCl, 150 mM NaCl, 5% glycerol, 0.01% IGEPAL, 10 mM imidazole) with 1x Halt Protease Inhibitor Cocktail, EDTA-Free (ThermoFisher), 40 mg of lysozyme, and 16 U Turbo DNase (Invitrogen). Cells were lysed with a BioSpec Tissue-Tearor (6 cycles of speed 6 then 4 cycles of speed 9 that were 30 s each with 30 s on ice between cycles). Lysates were cleared by centrifugation at 13,000 rpm for 15 minutes at 4°C. 8 mL His-Pur nickel–nitrilotriacetic acid resin 50% slurry (ThermoScientific) was added, and the NaCl concentration in the lysate was increased to 1 M before incubation for 60 minutes on at room temperature with end-to-end rotation. The resin was washed three times with 10 mL of 20 mM tris-HCl, 1 M NaCl, 5% glycerol, 0.01% IGEPAL, and 20 mM imidazole then eluted with 4 mL of 20 mM tris-HCl, 150 mM NaCl, 5% glycerol, 0.01% IGEPAL, 150 mM imidazole with 1x Halt Protease Inhibitor Cocktail, EDTA-Free (ThermoFisher) three times, on end-to-end rocker for 10 minutes each. Eluates were concentrated with Microcon 30,000 NMWL protein concentrators (MilliporeSigma) to a volume of about 300 µL. Samples were loaded onto 1 cm diameter, 38 ml Sephacryl S-200 High Resolution resin (GE Healthcare) size exclusion chromatography columns run with a BioLogic LP chromatography system (BioRad). A flow rate of 0.25 mL/minute was used, and 500 µL fractions were collected using SEC buffer (20 mM tris-HCl, 150 mM NaCl, 5% glycerol, 0.01% IGEPAL, 1 mM DTT, 1 mM EDTA). Fractions were combined, concentrated, and buffer exchanged using the same Microcon protein concentrators (Sigma) with 20 mM tris-HCL, 100 mM NaCl, 5% glycerol, 0.01% IGEPAL, and 0.1 mM DTT.

A truncated version of *M. smegmatis* RNase E containing amino acids 145-823 (deletion of part of the N-terminal intrinsically disordered region and deletion of the full C-terminal intrinsically disordered region) was used in this study as has been previously reported [2]. All RNase E variants used had N-terminal 6X his, 3X FLAG, tobacco etch virus (TEV) protease cleavage site, and 4X gly linker tags. A catalytically inactive version containing D694R and D738R mutations [17] was used as a control as described in [2]. RNase J had N-terminal 6X his, hemagglutinin (HA), tobacco etch virus (TEV) protease cleavage site, and 4X gly linker tags. A catalytically inactive version of RNase J containing D85K and H86A was also used as a control as described in [3].

### Overexpression and Purification of Recombinant

*M. tuberculosis* Proteins *M. tuberculosis* enzymes with C-terminal 6x his tags were overexpressed from pET plasmids in *E. coli*. Pellets were lysed in 20 mM Tris-HCl (pH 7.5), 500 mM NaCl, and 5% glycerol and loaded onto Ni-NTA resin using an AKTA purification system (Cytiva). Initial preps were washed with this buffer additionally containing 100 mM imidazole and 500 mM NaCl. High salt wash preps used 100 mM imidazole and 1 M NaCl. Proteins were eluted in a gradient from 75-500 mM imidazole then dialyzed in 50 mM Tris-HCl, pH 7.5, 100 mM NaCl, 5% glycerol, and 2 mM DTT buffer, run on a Superdex size exclusion column, and combined and concentrated.

### Preparation of RNA substrates

All RNA oligos used for this study are listed in Supplemental Table 3. Those used in Figure 1A-B, Supplemental Figure 1C, and Supplemental Figure 2C-F were purchased from Integrated DNA Technologies. In some cases, T4 polynucleotide kinase (New England Biolabs) was used to monophosphorylate 5’ ends of RNAs that were purchased with 5’ hydroxyls. Reactions were performed according to the instructions of the manufacturer and used directly in cleavage assays following heat inactivation of the kinase.

**Figure 1.**
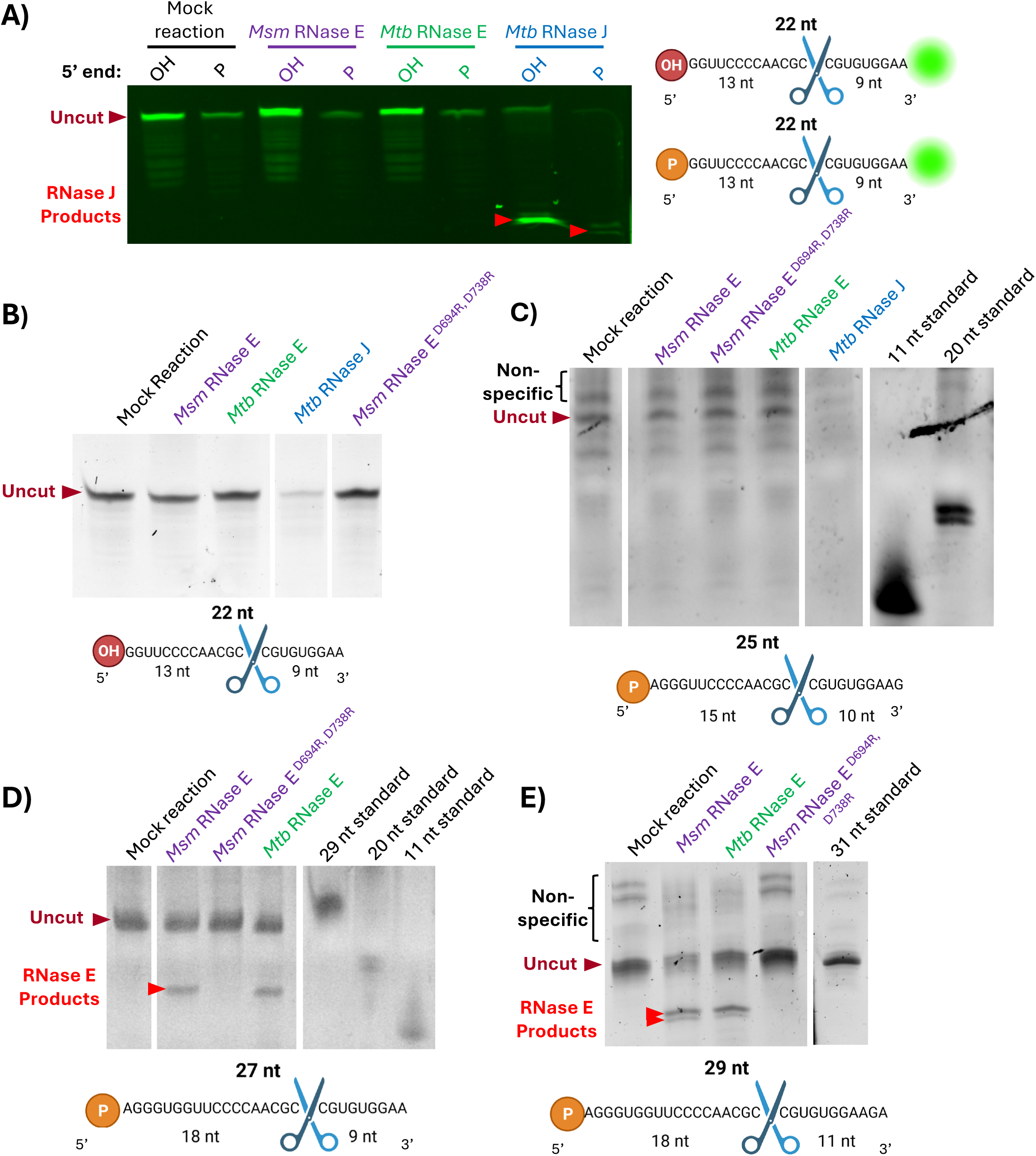
27 nt is the minimum length required for RNase E cleavage of an *atpB*-derived RNA sequence. All RNAs are derived from the *atpB* sequence of *M. smegmatis* and representative images of cleavage assays are shown. Gels are all 20% TBE urea-PAGE and were cropped to only show relevant lanes. RNase E ^D694R, D738R^ is expected to be catalytically inactive (Bandyra et al 2018). Nonspecific IVT products are noted. **A)** Cleavage of 1.5 ng of 22 nt 3’ FAM labeled oligo with 80 ng of enzyme. OH represents as purchased with a 5’ hydroxyl, and P represents PNK-treated to add a 5’ monophosphate. **B)** Cleavage of 100 ng of purchased 22 nt unlabeled oligo with 160 ng of enzyme. **C-E)** Cleavage assays with 80 ng of enzyme and RNAs made by IVT: **C)** of 40 ng of 25 nt RNA; **D)** 300 ng of 27 nt RNA; **E)** 300 ng of 29 nt RNA. The expected RNase E cleavage sites are indicated with scissors, and the sizes of the resulting expected cleavage products are shown.

The RNA substrates in Figure 1C-E, Figure 2, Figure 3, Figure 4, and Supplemental Figure 2 A-B were synthesized by *in vitro* transcription (IVT). dsDNA templates were made either by PCR for long RNAs or by annealing two primers together for short RNAs by adding 25 µM of each primer in 10 mM tris-HCl pH 7.9, 50 mM NaCl, and 1 mM EDTA in 20 µL reactions and heating at 95°C for 2 minutes then cooling 1.5°C per 1.5 minute cycle until 24.5°C.

**Figure 2.**
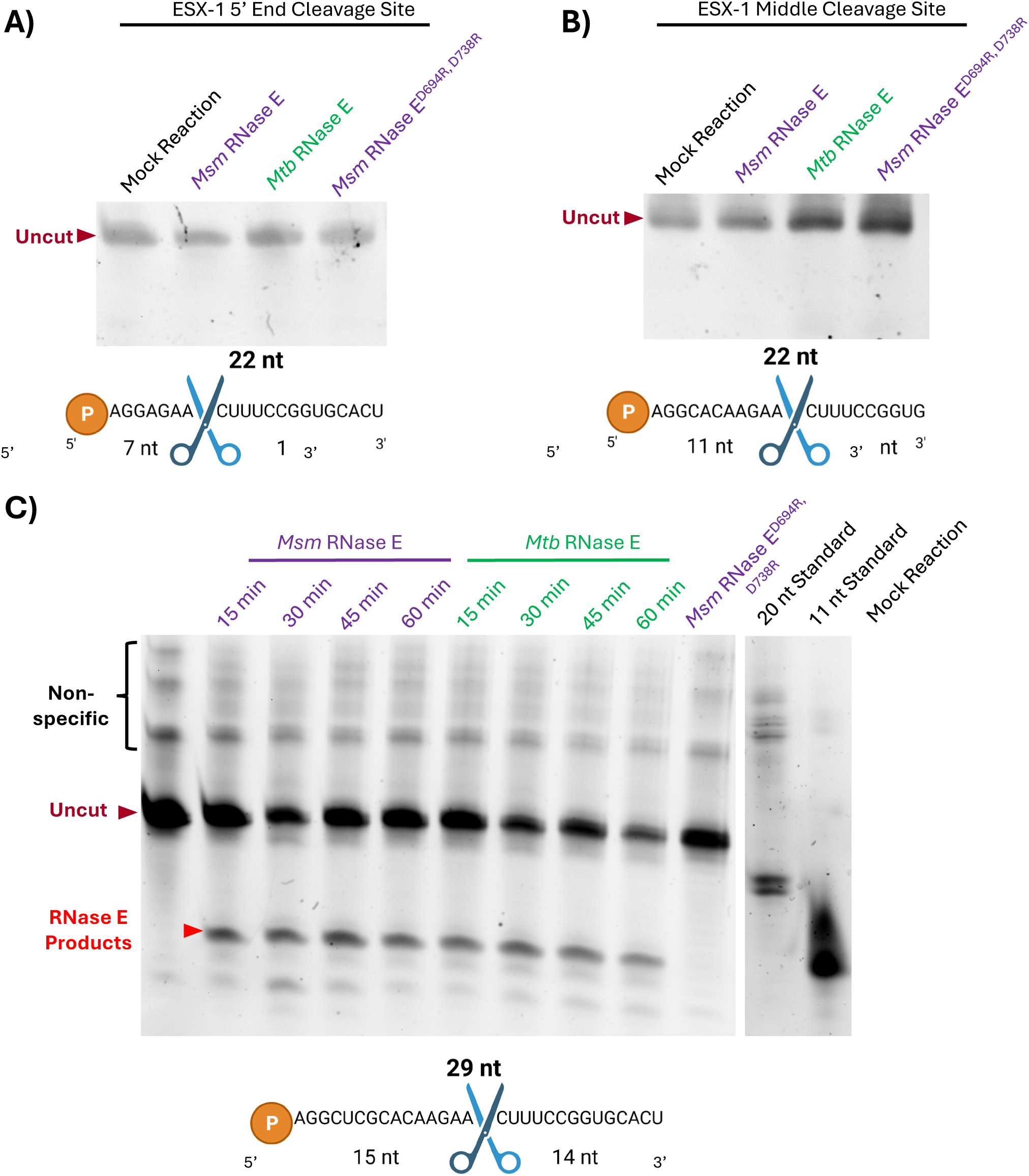
RNase E cleaves a 29 nt RNA sequence from the ESX-1 locus but not a 22 nt version. All RNAs are derived from intergenic region between *PPE68* and *esxB* in the ESX-1 locus of *M. smegmatis* and representative images of cleavage assays are shown. Gels are all 20% TBE urea-PAGE and were cropped to only show relevant lanes. Reactions contain 300 ng of RNA and 80 ng of enzymes. RNase E ^D694R, D738R^ is expected to be catalytically inactive (Bandyra et al 2018). Nonspecific IVT products are noted. Cleavage assay of a 22 nt oligo made by IVT with the cleavage site **A)** near the 5’ end and **B)** in the middle of the oligo. **C)** Timed cleavage assay with a 29 nt oligo containing an RNase E cleavage site in the middle. All RNAs had 5’ monophosphates. The expected RNase E cleavage sites are indicated with scissors, and the sizes of the resulting expected cleavage products are shown.

**Figure 3.**
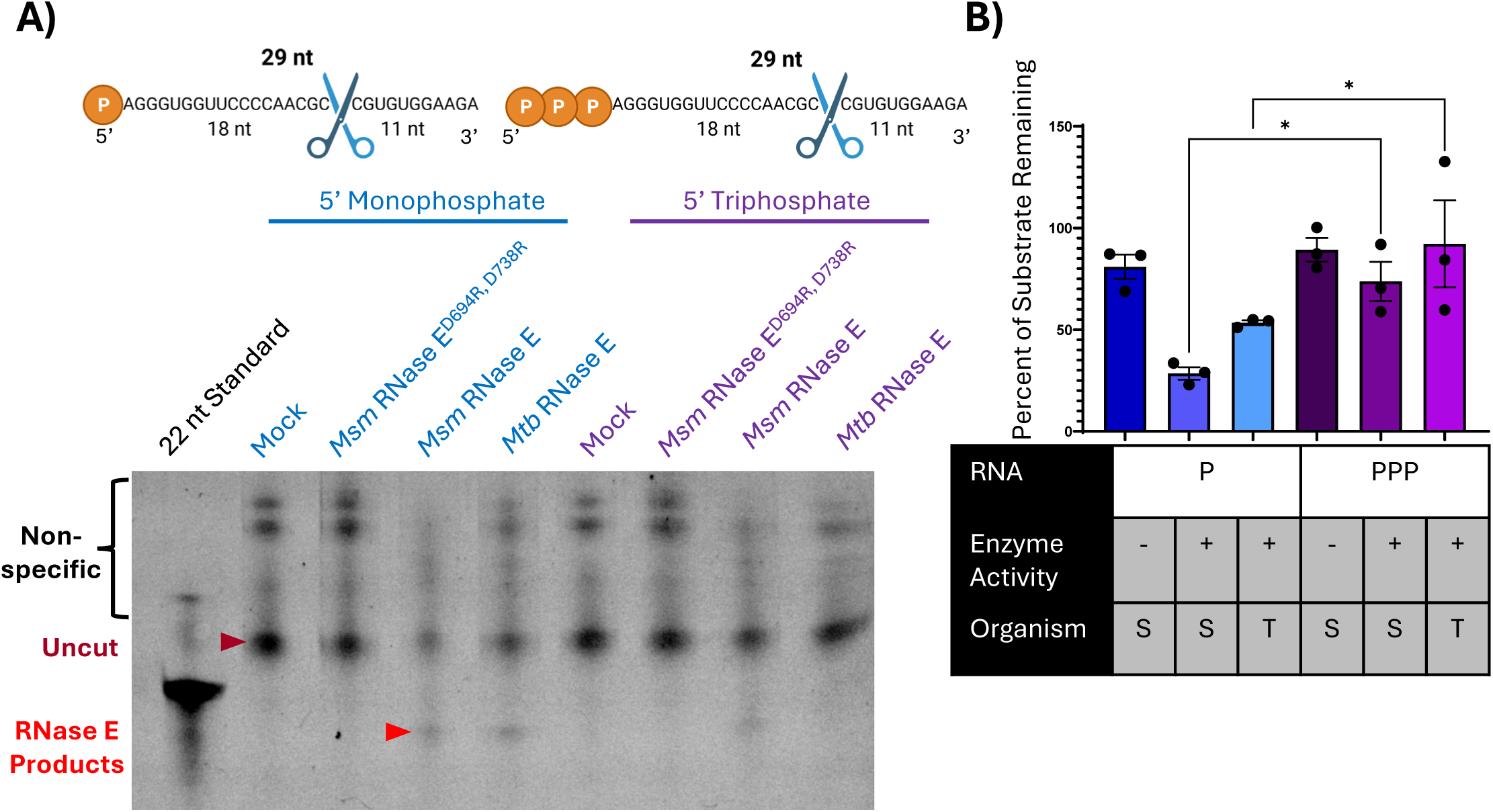
RNase E cleaves 5’ monophosphorylated substrates better than 5’ triphosphosphorylated substrates. Reactions contained 150 ng of RNA with 80 ng of enzymes. RNase E ^D694R, D738R^ is expected to be catalytically inactive (Bandyra et al 2018). Nonspecific IVT products are noted. **A)** Representative cleavage assay with the 29 nt *atpB* RNA made by IVT with a 5’ monophosphate or a 5’ triphosphate on a 20% TBE urea-PAGE gel. **B)** Quantification of three replicates of the experiment in A). The amount of substrate remaining for each condition was reported as a percentage of the mock reaction. In the RNA row, P represents 5’ monophosphorylated substrate, and PPP represents 5’ triphosphorylated substrate. In the enzyme activity row, (-) represents the catalytically inactive enzyme and (+) represents the catalytically active enzyme. In the organism row, S represents *M. smegmatis* RNase E and T represents *M. tuberculosis* RNase E. The bars represent the average of the three replicates, and error bars represent the standard error of the mean. An ordinary one-way ANOVA with Sidak’s multiple comparisons test was used for significance testing. * p ≤ 0.05

**Figure 4.**
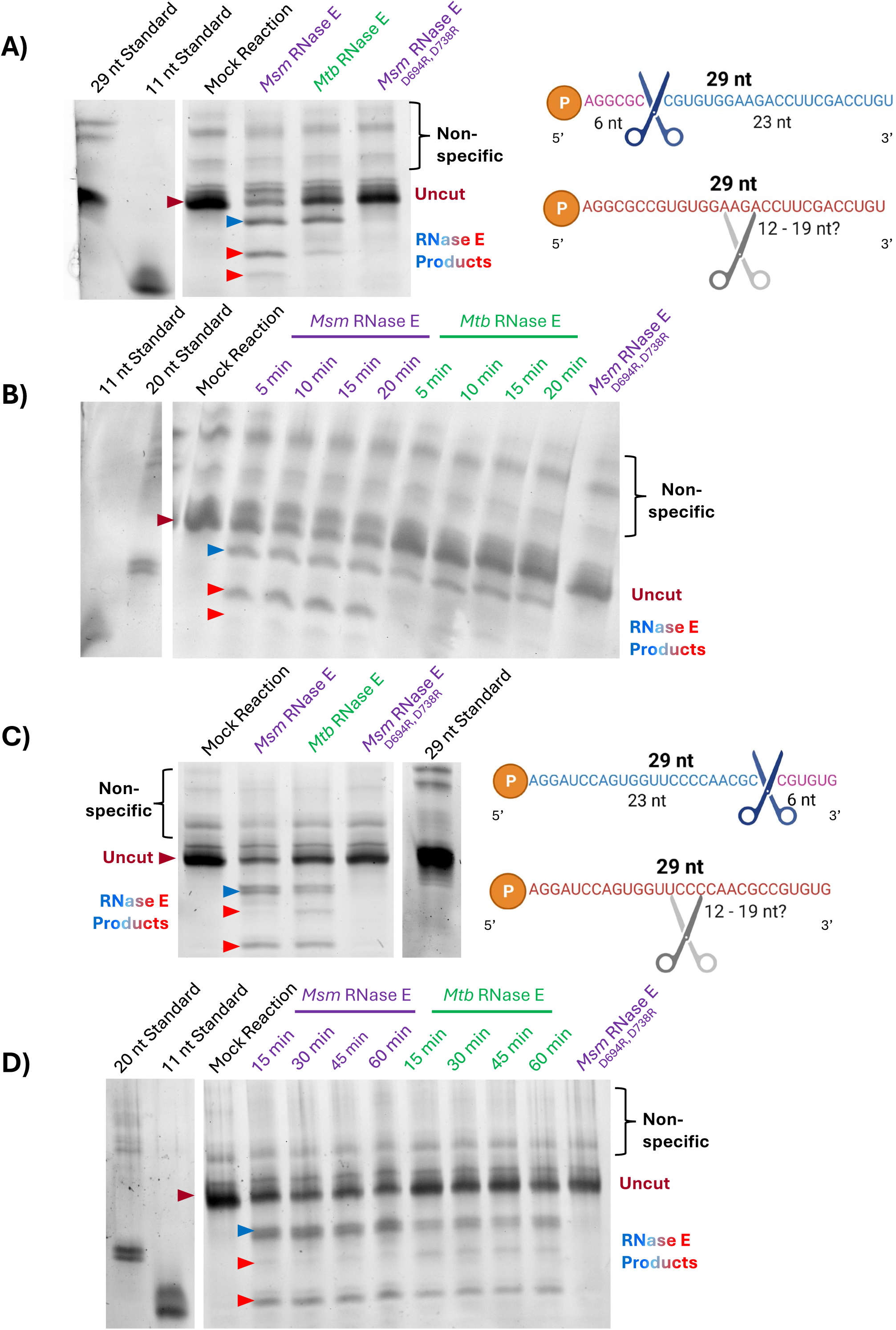
Both sequence and proximity to RNA ends influence RNase E cleavage positions. All RNAs are derived from the *atpB* sequence of *M. smegmatis* and representative images of replicated cleavage assays are shown. Gels were 20% TBE urea-PAGE and were cropped to only show relevant lanes. Reactions contained 300 ng of RNA with 80 ng of enzymes. RNase E ^D694R, D738R^ is expected to be catalytically inactive (Bandyra et al 2018). Nonspecific IVT products are noted. Cleavage of a 29 nt oligo made by IVT with the cleavage site motif located 6 nt from the 5’ end **A)** after one hour incubation and **B)** over a 20-minute time course. Cleavage of a 29 nt oligo made by IVT with the cleavage site motif located 6 nt from the 3’ end **C)** after one hour incubation and **D)** over a one-hour time course. The expected RNase E cleavage sites are indicated with blue scissors, and the resulting expected cleavage products are indicated with blue arrowheads on the gels. Gray scissors indicate the approximate positions of unknown observed cleavage events leading to two observed cleavage products with sizes in the range of 12-19 nt (red arrowheads on gels).

For generating RNAs with 5’ triphosphates (Figure 3 and Supplemental Figure 2A-B), HiScribe® T7 High Yield RNA Synthesis Kit (New England Biolabs) kit was used with a 20 µL reaction containing 7.5 µL of NTP + Buffer mix, 1.5 µL of T7 RNA Polymerase, and 250 ng of the dsDNA template and incubated overnight at 37°C. 2 µL of DNase I (New England Biolabs) and 30 µL of water were added to the reactions followed by incubation at 37°C for 15 minutes. RNA was purified using an RNA Clean and Concentrator-5 kit (Zymo) according to the manufacturer’s instructions with an elution volume of 11 µL.

RNAs with 5’ monophosphates were made in two different ways. To make the substrates used in Figure 3 and Supplemental Figure 2A-B, where we compared matched RNAs with 5’ triphosphates and 5’ monophosphates, we synthesized RNAs by IVT as described above and subsequently treated them with RNA 5’ Pyrophosphohydrolase (RppH) to convert 5’ triphosphates to 5’ monophosphates. RppH (New England Biolabs) was used according to the instructions from the manufacturer. Mock reactions with no RppH were set up in parallel, and the RNAs incubated in these mock reactions were used for direct comparisons of cleavage of monophosphorylated and triphosphorylated substrates. Following RppH treatment or mock treatment, RNA was purified using an RNA Clean and Concentrator-5 kit (Zymo) according to the manufacturer’s instructions with an elution volume of 11 µL.

The 5’ monophosphorylated RNAs used in Figure 1C-E, Figure 2, and Figure 4, which were not being directly compared to matched 5’ triphosphorylated RNAs, were synthesized with an excess of AMP to directly produce RNAs with largely monophosphorylated 5’ ends. HiScribe® T7 High Yield RNA Synthesis Kit (New England Biolabs) kit was used with a 20 µL reaction containing 7.5 µL of NTP + Buffer mix, 1.5 µL of T7 RNA Polymerase, 250 ng of the dsDNA template, 20 U of murine RNase inhibitor (New England Biolabs), 5 mM DTT, and 25 mM AMP and incubated overnight at 37°C. 2.5 U of Turbo DNase (Invitrogen), 1X Turbo DNase buffer, and 80 U murine RNase inhibitor (New England Biolabs) were added to the reactions, and these were brought to a final volume 100 µL and incubated at 37°C with 200 rpm shaking for 1 hour. RNA Clean and Concentrator-5 (Zymo) was used to purify RNA according to the manufacturer’s instructions with an elution volume of 11 µL.

### Cleavage Assays

Cleavage assays were done in 10 μL reactions in a buffer composed of 20 mM tris-HCl pH 7.9, 100 mM NaCl, 10 mM MgCl_2_, 10 μM ZnCl_2_, 0.5% glycerol, 0.01% IGEPAL, and 1 mM DTT. Reactions were incubated at 37°C for 1 hour unless otherwise indicated and were stopped by adding an equal volume of RNA loading dye (ThermoScientific) for unlabeled RNAs, or if the RNA was FAM-labeled, an equal volume of 12.5 mM EDTA pH 8.0 in formamide was used instead of loading dye. Reactions were then heated for 10 minutes at 70°C. Samples were run on TBE-urea PAGE gels. Those with unlabeled RNAs were stained with 40 mL of 1X TBE with 1X Sybr™ Gold (Invitrogen) on a rocker for 10 minutes then imaged with an Azure 600 imager at 302 nm excitation. FAM-labeled RNAs were imaged on an Azure 600 imager using the Cy3 channel (524 nm excitation). ImageJ with the FIJI plugin was used to quantify band intensities, which were normalized to a mock reaction with no enzymes where indicated. The amount of RNA and enzymes is noted in the figure legends.

### SDS-PAGE Gels

7.5% SDS-PAGE gels were stained with Bio-Safe Coomassie G-250 Stain (BioRad).

### Statistical Analysis

Statistical analysis was done with GraphPad Prism as indicated in individual figure legends.

## RESULTS

### 27 nt is the minimum length required for RNA cleavage by mycobacterial RNase E

We sought to investigate the cleavage preferences of mycobacterial RNase E using a simple *in vitro* system with a substrate containing a single cleavage site. We therefore purchased a 3’-6-carboxyfluorescein (FAM)-labeled 22 nt RNA oligo composed of sequence from the *atpB* gene, which we expected to contain exactly one RNase E cleavage site based on our previous work [2]. This oligo bore a 5’ hydroxyl group, and we used T4 polynucleotide kinase (PNK) to monophosphorylate a portion of it, as we expected RNase E to show a preference for monophosphorylated substrates [5, 7]. Reactions were performed with this oligo and full-length *M. tuberculosis* RNase E as well as *M. smegmatis* RNase E with truncations of portions of its intrinsically disordered regions. Previous work by us and others indicates that the intrinsically disordered region deletions are unlikely to impact catalytic activity on short substrates *in vitro* [2, 9, 17] . Surprisingly, we did not observe any cleavage of the 22 nt substrate by either *M. smegmatis* or *M. tuberculosis* RNase E, regardless of the 5’ end chemistry (Figure 1A). The substrate was cleaved by *M. tuberculosis* RNase J, which we included as a positive control for the assay given it has 5’ to 3’ exonuclease capability (Figure 1A). To test if the FAM label was interfering with the cleavage activity of RNase E, we repeated the assay with an unlabeled version of the same RNA, but we did not observe any cleavage by RNase E (Figure 1B). As we previously found that a 50 nt RNA including this 22 nt sequence was cleaved by RNase E [2], we then suspected that RNase E may have a minimum substrate length requirement. We used *in vitro* transcription (IVT) to synthesize variants of the substrate with progressively longer lengths. A 25 nt version was not cleaved by RNase E cleavage, but 27 nt and 29 nt versions showed cleavage (Figure 1C-E). This suggests that RNase E has a minimum length requirement of approximately 27 nt for the sequence tested here, which was a segment of the *atpB* gene.

To test if the minimum length requirement for RNase E cleavage was specific to the particular *atpB*-derived sequence tested above or applicable to other sequences, we tested an RNA sequence from the intergenic region between *PPE68* and *esxB* in the ESX-1 locus of *M. smegmatis,* which is known to contain an RNase E cleavage site [18]. We found that neither of two different 22 nt versions of the ESX-1 substrate with the cleavage site in different positions were cleaved by RNase E (Figure 2A-B). Increasing the length to 29 nt allowed for cleavage of the substrate by RNase E (Figure 2C). These results suggest that RNase E may have a minimum substrate length requirement that is unrelated to sequence. This result surprised us, as a previous report had shown that *M. tuberculosis* RNase E cleaved a 13 nt substrate [7].

### Recombinant mycobacterial RNase E and RNA helicase preparations can be contaminated with small amounts of highly active *E. coli* RNases

Preliminary experiments with an *M. smegmatis* RNase E mutant that was predicted to be catalytically inactive led us to suspect that some of our recombinant RNase E preparations were contaminated with *E. coli* RNases. To mitigate this, we increased the concentration of salt in the buffer used for wash steps in our nickel-nitrilotriacetic acid (Ni-NTA) affinity purification procedure (1 M NaCl rather than 500 mM NaCl). Although Coomassie-stained gels did not reveal major observable differences in purity of preparations done with the two wash protocols (Examples in Supplemental Figure 1A-B), we found that RNase E purified using the lower-salt wash protocol produced more cleavage products from a 29 nt RNA substrate (Supplemental Figure 1C). Furthermore, a preparation of the *M. tuberculosis* RNA helicase RhlE done with the lower-salt protocol produced cleavage products of the same sizes as the extra bands observed in the RNase E preps (Supplemental Figure 1C). As a helicase, RhlE is not expected to have any cleavage activity. These bands were not produced by RhlE purified using high-salt washes. These results suggest that both RNase E and RhlE purified from *E. coli* may co-purify with small amounts of *E. coli* RNases that could not be observed with Coomassie staining but nonetheless have detectable activity. High salt washing removed these contaminating activities and showed the expected results. With the exception of Supplemental Figure 1, all data shown in this work were generated using RNase E purified with the high-salt protocol.

### RNase E prefers to cleave 5’ monophosphorylated transcripts compared to 5’ triphosphosphorylated transcripts

The catalytic domain of RNase E in *E. coli* contains a pocket that can bind 5’ monophosphates of RNA substrates [5] which is required for efficient cleavage of many RNAs [19–21]. However, some RNA substrates are cleaved by RNase E regardless of 5’ end status suggesting that alternative 5’ end independent mechanisms for cleavage also exist [22, 23]. To assess the impact of 5’ end chemistry on cleavage by mycobacterial RNase E, we compared cleavage of substrates with 5’ monophosphates and 5’ triphosphates, as both are physiologically relevant. We synthesized a 29 nt *atpB* RNA by IVT, which we expected to be uniformly triphosphorylated. We then generated a monophosphorylated version by treatment of the IVT product with *E. coli* RppH and compared cleavage of this to the triphosphorylated version. We found that RNase E from both *M. smegmatis* and *M. tuberculosis* cleaved a greater proportion of the 5’ monophosphorylated substrate than the triphosphorylated substrate (Figure 3A-B), suggesting that it is stimulated by 5’ monophosphates similarly to *E. coli* RNase E. This is consistent with our recent report on the impact of the endogenous *M. tuberculosis* RppH on cleavage by *M. tuberculosis* RNase E [24].

### RNase J 5’ End Chemistry Preferences

We sought to use the substrates and assays developed here to additionally investigate the cleavage preferences of RNase J, a dual endonuclease and 5’ to 3’ exonuclease in mycobacteria [3]. *M. tuberculosis* RNase J was previously shown to have enhanced activity on a 5’ monophosphorylated versus a 5’ triphosphorylated RNA [15], and work on *M. smegmatis* and *Streptomyces coelicolor* RNase J suggested that 5’ monophosphates favor exonucleolytic cleavage while 5’ triphosphates favor endonucleolytic cleavage [3, 25]. We synthesized a longer 50 nt *atpB* RNA by IVT as previously reported [2] to increase the likelihood of including an RNase J endonuclease site and used RppH to generate monophosphates as described above. We found a clear RNase J cleavage preference for RNA with a 5’ monophosphate to RNA with 5’ triphosphate for both *M. smegmatis* and *M. tuberculosis* RNase J, but we did not observe any endonuclease products (Supplemental Figure 2A-B). To understand if 5’ hydroxyls on RNA substrates influence the cleavage mechanism of RNase J, we used PNK to generate a 5’ monophosphorylated version of the unlabeled 22 nt *atpB* RNA shown in Figure 1B. We found that *M. tuberculosis* RNase J cleaved the 5’ monophosphorylated substrate better than the to 5’ hydroxylated version (Supplemental Figure 2C-D), but *M. smegmatis* RNase J cleaved the two substrates similarly (Supplemental Figure 2E-F). We were unable to visualize any 1 nt products, so to confirm that the observed cleavage was due to 5’ to 3’ exonuclease activity, we purchased the same oligo sequence with a phosphorothioate bond between nucleotides 1 and 2 on the 5’ end. The phosphorothioate bond is expected to be cleaved more slowly than a phosphodiester bond, and therefore impact exonucleolytic cleavage but not endonucleolytic cleavage in our assay. We treated the oligo with PNK to ensure all versions of RNase J would have substantial cleavage abilities. This modified oligo showed significantly less cleavage with both *M. tuberculosis* and *M. smegmatis* RNase J (Supplemental Figure 2C-F), suggesting that we are observing exclusively exonuclease activity with these short oligos.

### Both sequence and proximity to ends appear to influence choice of cleavage position by RNase E

Mycobacterial RNase E preferentially cleaves in single-stranded regions between purines and cytidines [2]. The apparently strong preference for cleaving immediately upstream of cytidines contrasts with observations that proteobacterial RNase Es prefer to cleave in AU-rich regions [16, 26, 27] but aligns with mycobacteria being GC-rich organisms. However, we have consistently detected cleavage at some purine-cytidine dinucleotides in mycobacterial transcriptomes and not others [2, 28], indicating that further sequence preferences or other transcript features influence cleavage by RNase E. To investigate this, we assessed the impact of moving a verified RNase E cleavage site to different positions within a 29 nt substrate. We used sequences from the *atpB* gene and generated RNA substrates by IVT. We shifted the cleavage site assayed in Figure 1 to be 6 nt from either the 5’ end or the 3’ end to determine if RNase E could recognize the sequence of its original cleavage site in a different position, or if proximity to transcript ends also plays a role in cleavage site determination (Figure 4). We expected cleavage products of 6 nt and 23 nt from both of these new oligos if RNase E cut at the same sequence as in Figure 1. We were unable to detect the 6 nt product in any of our assays, possibly because this small product ran off the gel. However, we observed a band consistent with the expected 23 nt product when the cleavage site was located near the 5’ end of the RNA (Figure 4A-B). We also saw additional products, which were somewhere between 12-19 nt long based on our molecular weight standards, indicating that RNase E cut at one or more additional positions besides the expected cleavage site (Figure 4A-B). We found a similar result when the cleavage site was located near the 3’ end (Figures 4C-D), where we observed both an expected product and additional products. These results show that the sequence at the cleavage site itself is important for predicting cleavage, but the position within the RNA substrate is also important. These results also indicate that the failure of RNase E to cleave 22 nt and 25 nt substrates was not due to an absolute requirement for a minimum distance between the cleavage site and the 5’ or 3’ ends of the RNA.

### Possible product inhibition of RNase E

For several of the cleavage assays shown in this study, we stopped the cleavage reactions at various timepoints and assayed the products and remaining substrate (Figures 2D, 4B, and 4D). Interestingly, we never observed that the reactions went to completion. Moreover, there were no appreciable differences in product band intensity during the time intervals tested (5-20 minutes in Figure 4B, 15-60 minutes in Figures 2D and 4D). This suggests that RNase E may be subject to product inhibition, where cleavage products inhibit cleavage of additional substrate (see discussion).

## DISCUSSION

Here we have identified RNA features, including length and 5’ end chemistry, that impact RNase E cleavage *in vitro*. Consistent with expectations based on previous literature, we showed that both RNases E and J from both *M. smegmatis* and *M. tuberculosis* prefer 5’ monophosphorylated substrates to 5’ triphosphorylated ones. The catalytic domain of *E. coli* RNase E has a pocket that recognizes the 5’ end of the RNA [5]. Only a 5’ monophosphate, not a triphosphate or hydroxyl, can cause an allosteric effect to increase rates of catalysis when bound [5]. While no structure of mycobacterial RNase E has been reported, there is a high degree of sequence conservation of the catalytic domain with *E. coli* [6], so it is likely that a similar mechanism takes place to allow for more efficient cleavage of substrates with 5’ monophosphates. Interestingly, 5’ end chemistry has been shown to influence endonuclease versus exonuclease activity by RNase J. 5’ monophosphorylated substrates have been shown to promote exonuclease activity whereas 5’ triphosphorylated substrates are associated with endonuclease activity by RNase J from two Actinobacterial species: *Streptomyces coelicolor* [25] and *M. smegmatis* [3]. We observed that RNAs with 5’ monophosphates were cleaved more effectively than those with 5’ triphosphates by mycobacterial RNase Js, but we did not observe any products of endonucleolytic cleavage. A possible explanation for this observation is that no sites permissive for endonucleolytic cleavage by RNase J exist in the 22 nt and 50 nt *atpB* RNAs that we tested. Endonuclease sites remain elusive for RNase J in mycobacteria, so further investigations are needed to identify them before more can be learned about how RNase J decides between performing endonuclease or exonuclease activity.

We demonstrated that approximately 27 nt is the minimum RNA substrate length required for cleavage by mycobacterial RNase E. This contrasts with *E. coli* RNase E, which can cleave RNAs of 10 nt [19, 29] and 13 nt [19, 22, 29, 30]. RNase E from *E. coli* [8, 31] and *M. tuberculosis* [7] has been shown to form tetramers. In *E. coli*, 10 nt and 13 nt RNAs are bound by one tetramer whereas a 15 nt RNA is long enough to be shared by two tetramers [5]. In a crystal structure, the 15 nt RNA bound the 5’ end sensing pocket on one tetramer while simultaneously binding the active site on the second tetramer [5]. Shorter RNAs bound the 5’ end sensing pocket and active site of adjacent protomers within a single tetramer [5]. Thus, RNA length may influence the mode of binding by RNase E. Additionally, a report showed that an RNase E mutant that forms more multimers was able to more efficiently cleave long but not short RNAs in *E. coli* [30]. This suggests that RNase E multimerization state may contribute to selecting substrates for degradation by length. However, it is unclear why the minimum length requirement for mycobacterial RNase E differs from that of *E. coli*; a structure of mycobacterial RNase E bound to a substrate will likely be required to answer this question.

Our finding that substrates shorter than 27 nt were not cleaved by mycobacterial RNase stands in contrast to a previous report with *M. tuberculosis* RNase E in which a 13 nt oligo was cleaved [7]. Since we did not test the specific RNA sequence used in that report, it is possible that the different results are related to use of different substrate sequences. However, it should be noted that it is also possible that the RNase E preparations used in [7] were contaminated with co-purifying *E. coli* RNases. We were surprised to find contamination with *E. coli* RNases in our recombinant protein preparations despite them appearing pure on Coomassie-stained gels. Purifying catalytically inactive mutants helped us to identify this contamination and assess the effectiveness of the purification protocol changes that we made to eliminate the contamination. Specifically, including 1 M NaCl in the Ni-NTA affinity resin wash buffer effectively removed the contaminating activity from both RNase E and RhlE (RNA helicase) preparations. Mycobacterial RNase E contains two intrinsically disordered regions [6] and RhlE is also predicted to have a long intrinsically disordered region; these could potentially contribute to the binding of other *E. coli* proteins. Based on our experience, we highly recommend using high salt washing and purifying catalytically inactive enzymes in parallel with active protein preparations when recombinantly purifying mycobacterial RNase E and RhlE from *E. coli* to ensure high purity.

In organisms where cleavage patterns of RNase E have been studied globally, no consensus sequence has been identified, which is perhaps intuitively consistent with the requirements of an RNase with a major role in bulk mRNA degradation. In *E. coli* and *Salmonella typhimurium,* which have genomes that are approximately 51-52% G+C, RNase E preferentially cleaves when an uracil is in the +2 position [16, 26, 27]. In GC-rich *M. smegmatis* (∼67% G+C), RNase E preferentially cleaves 5’ of cytidines [2]. We show here that a known cleavage site sequence was still cut when moved to positions near the 5’ or 3’ end of a short substrate, but additional products were also made, showing that RNase E cleaved at additional positions nearer to the center of the substrate. This adds complexity to predicting cleavage positions for RNase E because both sequence and context matter. This result is consistent with a recent study in which we showed that mutating the cytidine at a major RNase E cleavage site in the ESX-1 locus (studied here in Figure 2) did not prevent cleavage at or near that position in *M. smegmatis* cells, indicating that other sequence, secondary structure, or positional features were more important for specifying cleavage at that location [18].

We observed that RNase E cleavage reactions rarely went to completion *in vitro,* suggesting that RNase E may be inhibited by its own cleavage products. It has been proposed that cleavage products may remain tightly associated with *E. coli* RNase E to prevent cleavage of additional substrates *in vitro* [32] and an autoinhibitory motif has been identified that was speculated to suppresses enzyme activity by slowing the release of cleavage products [17]. Our data in mycobacteria are consistent with this model. While beyond the scope of this study, more evidence is needed to define the extent to which mycobacterial RNase E binds its own cleavage products to more concretely determine if and how it is inhibited by its own products.

Finally, an interesting note is the remarkable functional similarity between *M. smegmatis* and *M. tuberculosis* RNase E. These two proteins consistently showed the same cleavage site positions, 5’ end preferences, length requirements, sequence context importance, and possible product inhibition properties. This supports the value of *M. smegmatis* as a comparable nonpathogenic model for studying *M. tuberculosis* RNA metabolism and the roles of RNase E in particular.

## ACKNOWLEDGEMENTS

We thank members of the Shell lab for thoughtful discussions and suggestions.

## CONTRIBUTIONS

**Abigail R. Rapiejko:** conceptualization, methodology, formal analysis, investigation, writing – original draft, data visualization. **Manchi Reddy:** investigation. **James C. Sacchettini:** methodology, supervision, funding acquisition. **Scarlet S. Shell:** conceptualization, writing – review & editing, supervision, project administration, funding acquisition.

## SUPPLEMENTARY DATA

**Supplemental Figures 1-2**

**Supplemental Table 1. Gene IDs**

**Supplemental Table 2. Expression Plasmids**

**Supplemental Table 3. RNA oligos**

## CONFLICT OF INTEREST

None

## FUNDING

This work was supported in part by NIH-NIAID award 5TP01AI143575-02 to SSS and JCS, by NIH-NIAID award R21 AI156415-01A1 to SSS, by NSF-CAREER award 1652756 from the Directorate of Biological Sciences to SSS, an award from the Potts Foundation to SSS, DoEd GAANN training grant P200A240115 to ARR, and a Dr. Armand P. Ferro and Mary H. Ferro Summer Fellowship to ARR. The funders had no role in study design, data collection and analysis, decision to publish, or preparation of the manuscript.

---

**Supplemental Figure 1.**
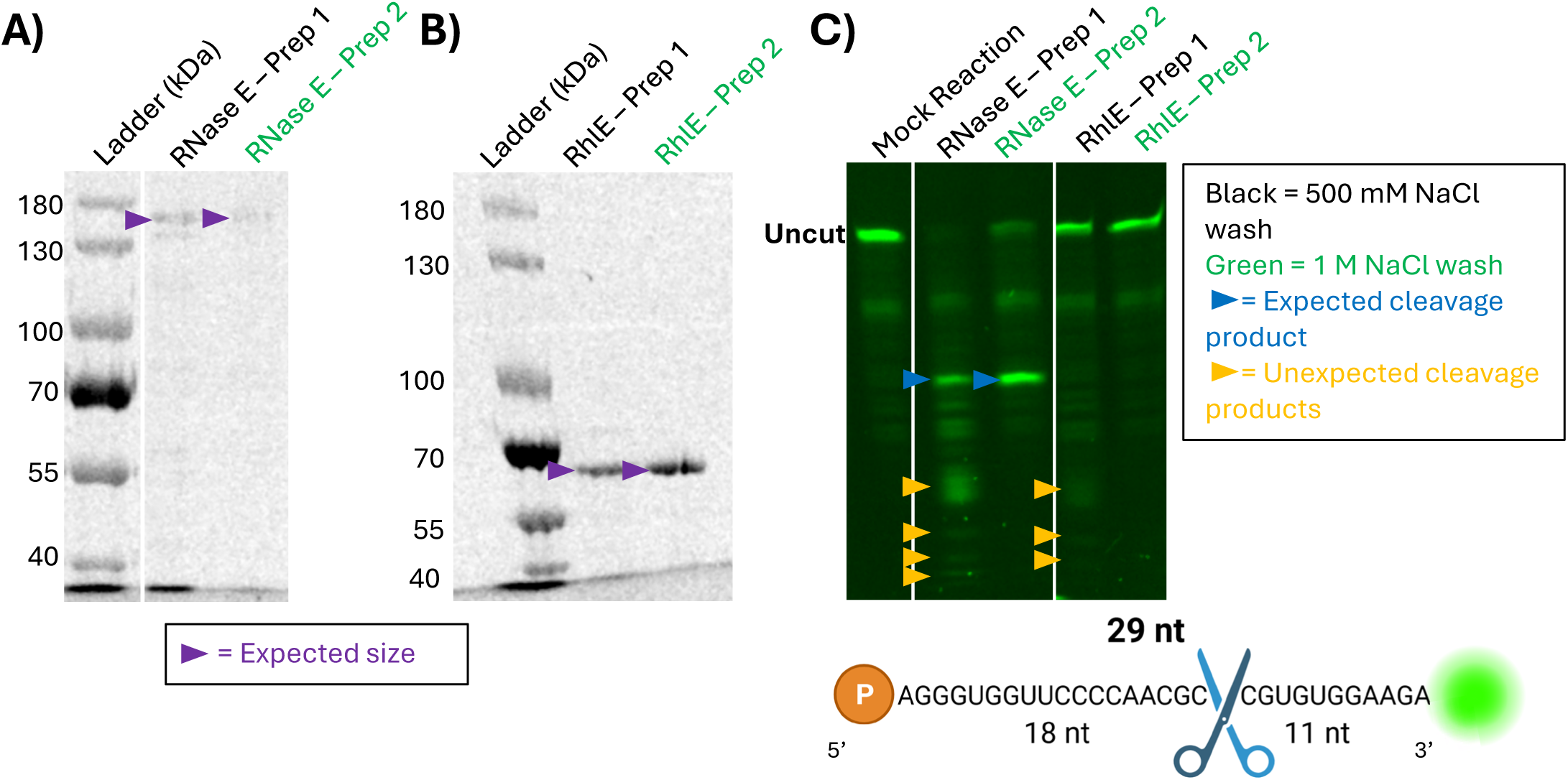
Mycobacterial RNase E can be contaminated with small amounts of *E. coli* RNases. All RNAs are derived from the *atpB* sequence of *M. smegmatis* and representative images of cleavage assays are shown. Gels were cropped to only show relevant lanes. Coomassie stained 7.5% SDS-PAGE gels of preps of *M. tuberculosis* **A)** RNase E and **B)** RhlE. High salt washing during IMAC purification is indicated by the green font, and the purple arrow indicates the expected band size. **C)** Cleavage assay with 20 ng of RNA and 80 ng of enzymes run on a 20% TBE urea-PAGE gel of the preps from A) and B) with a purchased 29 nt 3’ FAM labeled RNA. Blue arrows indicate the expected RNase E cleavage product and yellow arrows indicate extraneous bands due to contaminating RNases. Blue scissors indicate the position of the RNase E cleavage site mapped in Zhou et al 2023.

**Supplemental Figure 2.**
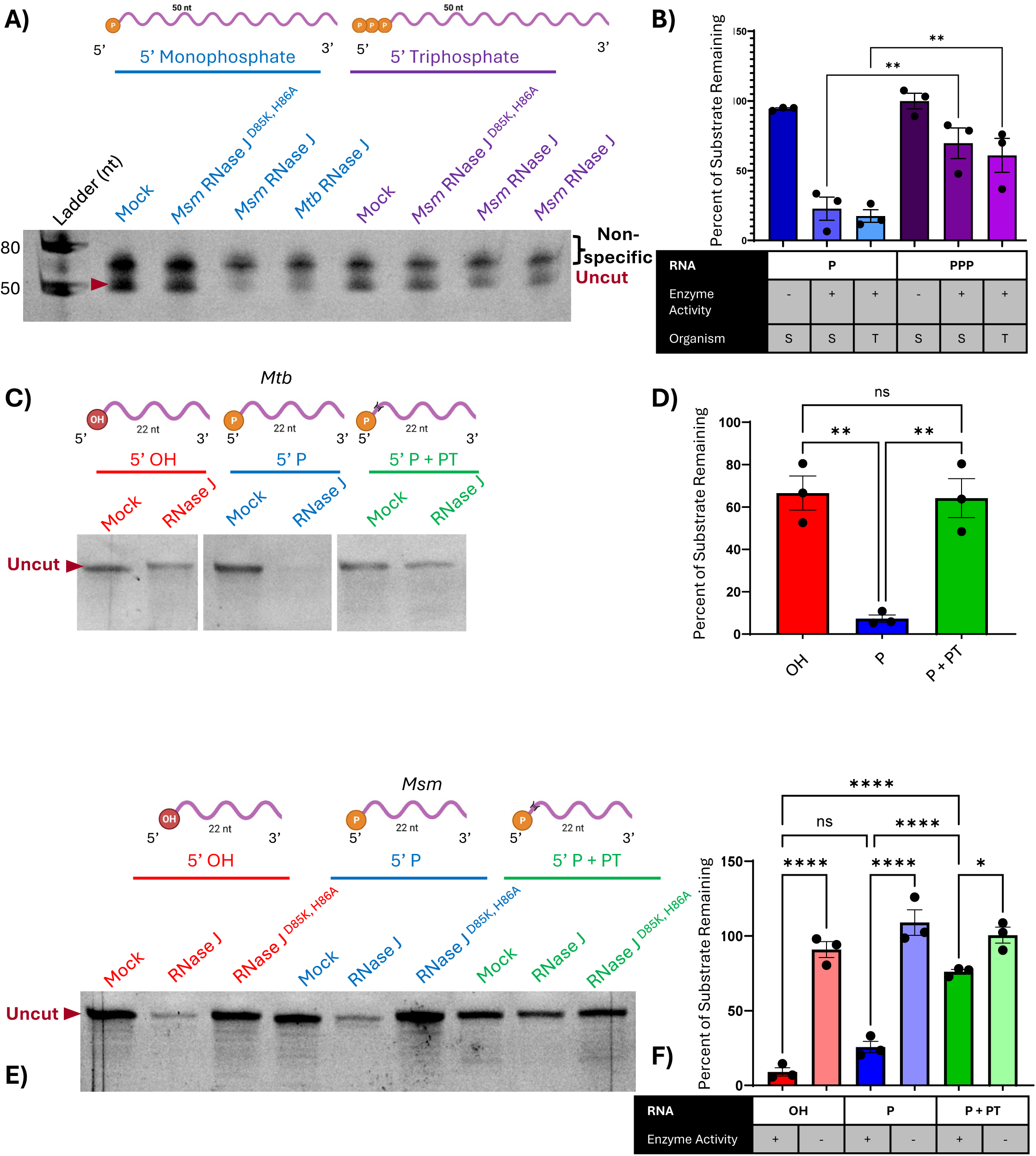
The impact of 5’ end chemistry on cleavage by mycobacterial RNase J. All RNAs are derived from the *atpB* gene of *M. smegmatis* and representative images of cleavage assays are shown. Gels were cropped to only show relevant lanes. Reactions with 40 ng of RNA and 80 ng of enzyme were used. RNase J ^D85K, H86A^ is expected to be catalytically inactive (Taverniti et al 2011). Nonspecific IVT products are noted. **A)** Cleavage assay of 50 nt RNAs made by IVT with RNase J run on a 15% TBE urea-PAGE gel, quantified in **B)**. **C-F)** Cleavage assays using a purchased 22 nt oligo with a 5’ hydroxyl, 5’ monophosphate, or a 5’ monophosphate with a phosphorothioate bond between positions 1 and 2 on the 5’ end. RNAs were run on 20% TBE urea-PAGE gels. **C)** *M. tuberculosis* RNase J quantified in **D). E)** *M. smegmatis* RNase J quantified in **F)** . The amount of substrate remaining for each condition was reported as a percentage of the mock reaction for three replicates. OH represents 5’ hydroxyl, P represents 5’ monophosphate, and P + PT represents 5’ monophosphate with the phosphorothioate modification. In the enzyme activity row, (-) represents the catalytically inactive enzyme and (+) represents the catalytically active enzyme. In the organism row of B), S indicates *M. smegmatis* RNase E and T indicates *M. tuberculosis* RNase E. The bars represent the average of the 3 replicates, and error bars represent the standard error of the mean. An ordinary one-way ANOVA with Sidak’s multiple comparisons test was used for significance testing. ns p > 0.05, * p ≤ 0.05, ** p ≤ 0.01, **** p ≤ 0.0001

## Notes

### Competing Interest Statement

The authors have declared no competing interest.

